# CUE: CpG impUtation Ensemble for DNA Methylation Levels Across the Human Methylation450 (HM450) and EPIC (HM850) BeadChip Platforms

**DOI:** 10.1101/2020.05.30.107094

**Authors:** Gang Li, Laura Raffield, Mark Logue, Mark W Miller, Hudson P. Santos, T.Michael O’Shea, Rebecca C. Fry, Yun Li

## Abstract

DNA methylation at CpG dinucleotides is one of the most extensively studied epigenetic marks. With technological advancements, geneticists can profile DNA methylation with multiple reliable approaches. However, profiling platforms can differ substantially in the CpGs they assess, consequently hindering integrated analysis across platforms. Here, we present CpG impUtation Ensemble (CUE), which leverages multiple classical statistical and modern machine learning methods, to impute from the Illumina HumanMethylation450 (HM450) BeadChip to the Illumina HumanMethylationEPIC (HM850) BeadChip. Data were analyzed from two population cohorts with methylation measured both by HM450 and HM850: the Extremely Low Gestational Age Newborns (ELGAN) study (*n*=127, placenta) and the VA Boston Posttraumatic Stress Disorder (PTSD) genetics repository (*n*=144, whole blood). Cross-validation results show that CUE achieves the lowest predicted root mean square error (RMSE) (0.026 in PTSD) and the highest accuracy (99.97% in PTSD) compared with five individual methods tested, including k-nearest-neighbors, logistic regression, penalized functional regression, random forest and XGBoost. Finally, among all 339,033 HM850-only CpG sites shared between ELGAN and PTSD, CUE successfully imputed 289,604 (85.4%) sites, where success was defined as RMSE < 0.05 and accuracy >95% in PTSD. In summary, CUE is a valuable tool for imputing CpG methylation from the HM450 to HM850 platform.

## Introduction

DNA methylation of cytosine residues at CpG dinucleotides is one of the most extensively studied epigenetic marks. Recent literature provides evidence regarding its important role not only in normal development but also in risk and progression of many diseases ^1-8^. A wide range of biological processes are dependent on DNA methylation status, including gene transcription, X-chromosome inactivation, cell differentiation, cancer progression and other critical life events or processes such as aging ^4, 7^. Therefore, studies of DNA methylation are of great interest and importance but present challenges for a number of reasons including but not limited to the following: 1) DNA methylation levels can be dynamic, varying over time, across different environments, developmental stages, and tissues or cell lines; 2) correlation of methylation levels between CpG sites decreases dramatically with distance, for example with correlation coefficients typically < 0.5 when two CpG sites are merely >500bp apart; and 3) there are multiple methods for the measurement of DNA methylation (see section 2 for a more detailed review).

With the emergence of powerful technologies such as DNA methylation arrays ^9^ and bisulfite sequencing, geneticists are able to profile DNA methylation levels at increasingly higher resolutions. Methylation microarrays have become very popular due to lower cost and higher throughput than bisulfite sequencing approaches. However, new platforms assaying an increasing number of CpG sites have replaced old ones every few years ^10-12^. Different platforms (for example, the widely used Illumina HumanMethylation27, HumanMethylation450, and MethylationEPIC BeadChips) target different CpG sites across the genome and have different marker densities. In addition, different biochemical or experimental techniques can be used to quantify methylation levels (e.g., type-I versus type-II assays adopted by the Illumina methylation arrays), further hindering joint analysis of data from multiple platforms.

Two aforementioned microarrays, the Illumina HumanMethylationEPIC (HM850) and HumanMethylation450 (HM450) BeadChips, are the most commonly used microarrays to measure DNA methylation levels. While HM450 investigates 485,577 probes spanning 96% of CpG islands and 92% of CpG shores across a moderate number of genes ^9^, HM850 provides much more comprehensive coverage with the additional 413,743 CpG sites located farther outside CpG islands. As new arrays are introduced on a fairly regular basis, and as old chips have typically not been available (HM450 was discontinued) for purchase once the new chips have been introduced, researchers will increasingly encounter data generated from a combination of different arrays. Such data have largely constrained pooled analysis, where investigators typically focus on the probes shared between the two platforms ^13-15^. This is a prudent and convenient approach which does not necessitate reevaluation of all samples, for example using an updated HM850 array, which would be time consuming, expensive and wasteful of valuable tissue samples. However, applying such an approach to a dataset comprised of a mix of HM450 and HM850 implies an unfortunate waste of HM850 data where more than 40% of data will not be used in pooled analyses. In this study, we present the CpG impUtation Ensemble (CUE), an ensemble learning framework which leverages several machine learning algorithms and traditional statistical models to efficiently integrate data from these two platforms. While several existing methods designed for imputing sequencing-density methylation levels require hundreds of genomic features (e.g., those from the ENCODE project ^16^) to impute each missing CpG methylation site, the present study highlights a relatively simple and more widely-applicable imputation regime which only requires methylation measurements from the HM450 BeadChip.

Because of the practical needs for imputation in the context of DNA methylation data as well as the success of imputation methods in other genetic settings ^17-19^, a number of DNA methylation imputation methods have been proposed. Among them, support vector machines (SVM) and a hybrid of SVM and other models predominate in the DNA methylation imputation literature ^20-27^. Many of these methods assume that methylation status is binary. In other words, a CpG site is either methylated or unmethylated for an individual and thus imputation becomes a classification problem. Almost all methods proposed prior to 2014 predicted the average methylation status for genomic regions, where each region encompasses multiple CpG sites ^28^. All of the studies reported accuracy exceeding 90%.

Dichotomizing methylation status, as adopted by previously published SVM based methods, can lead to the loss of biologically meaningful information carried by intermediate raw beta values (*β*). The beta value is the ratio of intensities between methylated and unmethylated alleles and is one standard quantitative measure of DNA methylation levels. *β* values range from 0 to 1 with 0 being completely unmethylated and 1 completely methylated. With advances in data science, particularly in areas of machine learning and deep learning ^29^, several algorithms have been successfully employed for methylation imputation and reported to outperform the earlier SVM based methods. For example, Zhang *et al*. ^28^ in 2015 employed a random forest (RF) classifier to predict methylation levels with five groups of features selected from the ENCODE Project, achieving 96% accuracy. Angermueller *et al*. ^30^ in 2017 adopted a deep learning method to provide an accurate prediction of single-cell DNA methylation states, which achieved performances similar to the previous SVM or RF based methods. In the same year, BoostMe ^31^, based on the state-of-the-art boosting algorithm XGBoost, achieved the same level of accuracy as RF, but with increased computational efficiency.

In the present study, we set out to impute data for the HM850 array using data from the HM450, for increased coverage of the epigenomic landscape. We have developed a general CUE framework and compared it against five available methods to assessed their performance to impute data. The five methods evaluated were k-nearest-neighbors (KNN), logistic regression (Logistic) and penalized functional regression (PFR) model ^32, 33^, random forest (RF) and XGBoost. We applied the CUE framework to imputation in two cohorts with methylation measured both by HM450 and HM850: the VA Boston Posttraumatic Stress Disorder (PTSD) genetics repository ^34^ (144 whole blood samples) and the Extremely Low Gestational Age Newborns (ELGAN) study ^35^ (127 placenta samples). We subsequently examined the imputation results and filtered out the low-quality probes. Accurately imputed methylation values could subsequently improve power in downstream analysis, for example for associating methylation profiles with phenotypic traits of interest, widely referred to as epigenome-wide association studies (EWAS). Pre-trained imputation models for placenta or whole blood samples can be found on https://github.com/GangLiTarheel/CUE, as well as code for applying CUE to new reference datasets assessed using multiple methylation arrays.

## Results

In this study, DNA methylation data were used from both the HM450 and HM850 platforms derived from two human cohorts, namely the VA Boston Posttraumatic Stress Disorder (PTSD) genetics repository ^34^ (144 whole blood samples) and the Extremely Low Gestational Age Newborns (ELGAN) study ^35^ (127 placenta samples). Five methods were assessed comprehensively for their ability to impute data for the HM850 platform using data from the HM450: three traditional statistical methods, k-nearest-neighbors (KNN), logistic regression (Logistic) and penalized functional regression (PFR) model ^32, 33^; and two modern machine learning algorithms: random forest (RF) and XGBoost. Method performance was systematically evaluated using six-fold cross-validation on the two cohorts (ELGAN and PTSD) separately.

### Cross Validation Results on ELGAN and PTSD

In this paper, we focused on imputation within tissue type for two reasons. First, for most studies, samples are usually collected for the same tissue. Second, differences in methylation patterns across tissue types prevent accurate imputation across tissues. Imputation quality was assessed within each cohort by conducting six-fold cross validation, separately on placenta samples from the ELGAN study and whole blood samples from the PTSD study.

Note the time complexity of the method is O(*n*) and thus each target probe can easily impute in parallel to decrease clock computation time, where *n* is the number of the HM850-specific probes (in this case *n*=339,014 HM850-specific probes).

For the ELGAN placenta dataset, RF achieved the smallest root mean square error (RMSE) (0.099) and the highest accuracy (measured by dichotomizing DNA methylation level at a cutoff of 0.5) (94.60%) among the five imputation tools that were compared (**Table 1**). KNN, PFR and XGBoost performed less well than RF with regard to RMSE (decreases by 0.004-0.025), and had 0.34%-1.96% loss in terms of classification accuracy for dichotomous methylation status. Logistic regression was the fastest algorithm and achieved an accuracy higher than 90% but performed less well in RMSE.

**Table 1.** Imputation Performances in the ELGAN and PTSD datasets. For all the computational results reported in tables of this paper, 15 CPUs were used. Logistic regression did not converge (D.N.C.) for the PTSD dataset. The fastest method computation time and the highest method accuracy are displayed in bold for each dataset.

|  | **ELGAN Performances** |  |  | **PTSD Performances** |  |  |
| --- | --- | --- | --- | --- | --- | --- |
|  | Accuracy | RMSE | Time | Accuracy | RMSE | Time |
| **KNN** | 92.64% | 0.124 | 2.5hrs | 98.02% | 0.054 | **3hrs** |
| **Logistic** | 91.76% | 0.263 | **2.5hrs** | D.N.C. | D.N.C. | D.N.C. |
| **PFR** | 93.50% | 0.114 | 6hrs | 98.41% | 0.044 | 18.5hrs |
| **RF** | **94.60%** | **0.099** | 3hrs | 98.02% | 0.054 | 5hrs |
| **XGBoost** | 94.26% | 0.103 | 3hrs | **98.59%** | **0.040** | **3hrs** |

Similarly, six-fold cross validation results were obtained in the PTSD dataset. Among the five single imputation results, XGBoost outperformed the other methods, in contrast to RF for the ELGAN dataset. Specifically, XGBoost achieved the fastest speed (3 hrs), the smallest RMSE (0.04) and the highest accuracy (98.59%) (**Table 1**). In fact, RF was not even the second-best performing method. The PFR approach achieved a 0.39% higher accuracy and a lower (by 0.01) RMSE than RF. Compared with the results from the ELGAN dataset, these results suggest that there is no uniformly best single imputation method across different tissues or datasets, which inspired the development of the ensemble imputation framework.

We further reported at each CpG site which method outperformed all other single imputation methods, then showed the proportion of CpG sites for which each method performed best (**Figure 1**). Results showed that all four viable imputation methods (we excluded logistic regression since it failed to converge) perform best at some CpG sites, which again motivates the development of an ensemble imputation framework. For more than 42% of the CpG sites, RF achieved the lowest RMSE (**Figure 1**). In contrast, PFR was the best imputation model for 26% of the CpG sites while XGBoost was the best for ∼30% CpG sites (**Figure 1**). Lastly, for 1.7% of the probes (5,596 probes) KNN performed the best (**Figure 1**). In total, RF and XGBoost outperformed the other methods for ∼70% of CpG sites.

**Figure 1.**
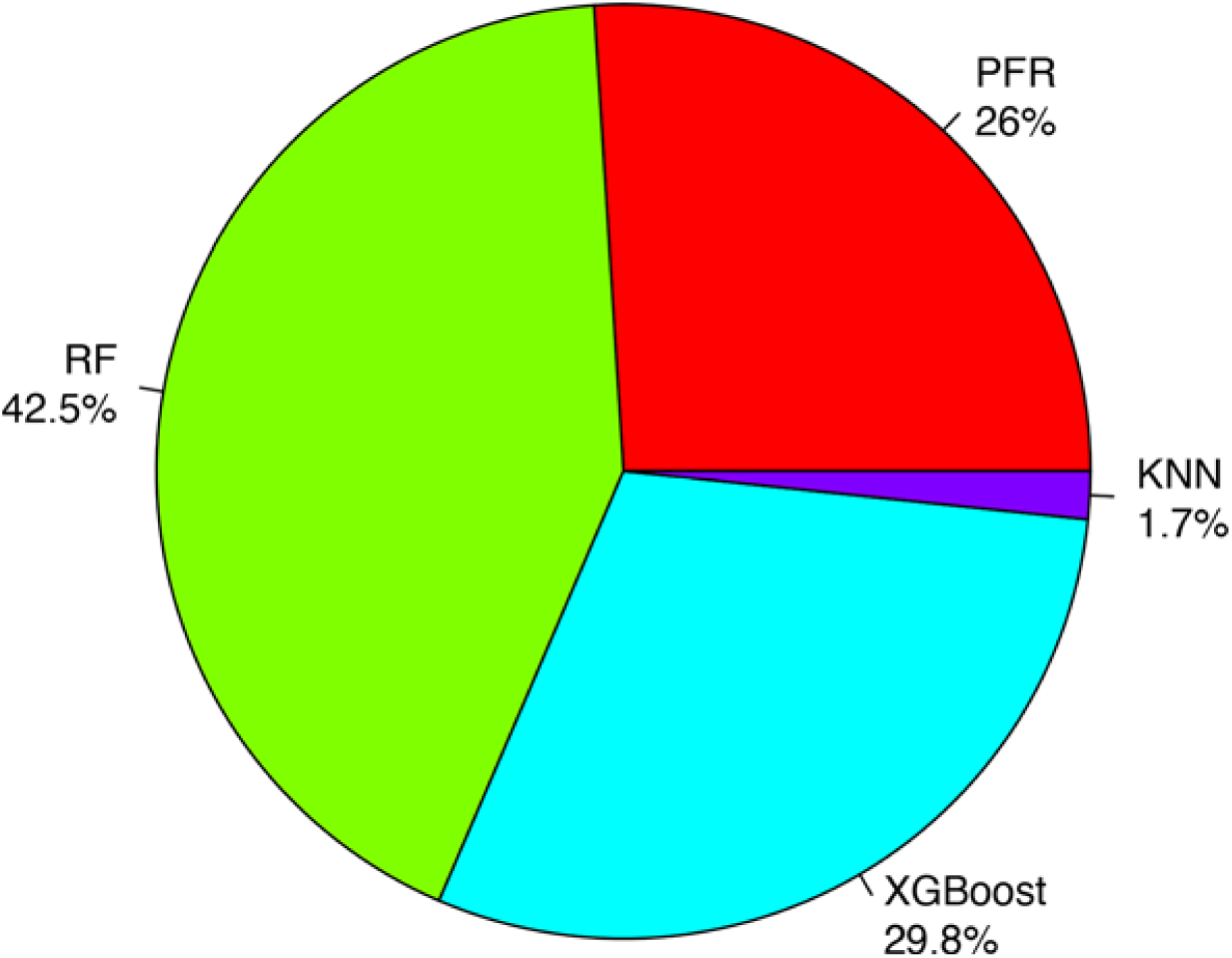
Proportions of CpG Sites where Each Method Wins in PTSD.

### Ensemble Imputation Results with CUE within the PTSD dataset

Based on these inconsistent results in terms of which method had the best performance, CUE was developed to improve the prediction of methylation values at HM850 specific CpG sites. Probes were filtered for those that failed to pass quality control (QC) criteria (RMSE <0.05 and accuracy >95% at CpG site level). Prior to QC, all the methods predicted RMSE below 0.06 and CUE showed the best performance (**Figure 2**). The predicted RMSE of the individual tools can be reduced by 5.8%-30.3% with CUE. After post-imputation QC, the predicted RMSE of all the tools were reduced by 37.9%-50.0%. Among the 339,033 HM850-only CpG sites shared between ELGAN and PTSD, CUE out-performed the individual methods at 289,604 (85.4%) sites. Specifically, CUE achieved the lowest predicted RMSE (0.026) and the highest accuracy (99.97%), compared with individual methods which had RMSE ranging 0.029-0.036 (improved by 10.0%-27.4%) and with accuracy 99.95%-99.97%.

**Figure 2.**
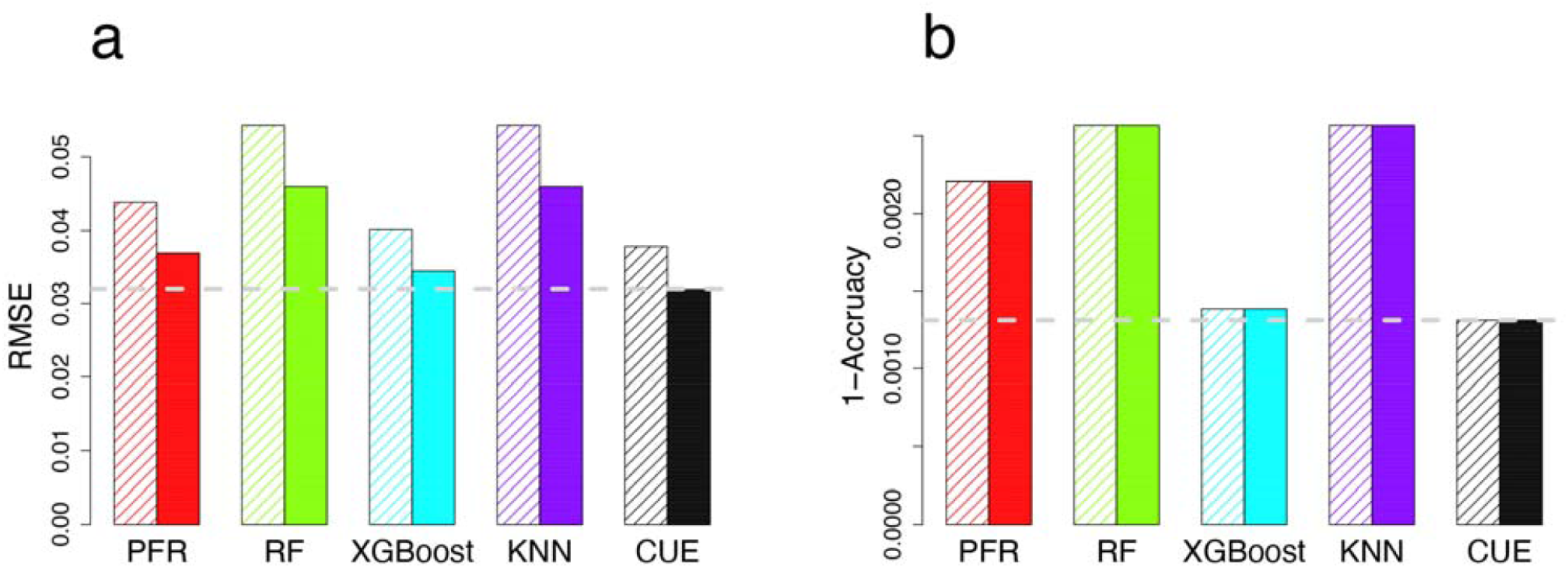
Imputed performances for probes before (left, hatched) and after QC (right, no hatches) in PTSD. (**a**) RMSE comparisons. (**b**) 1-Accuracy (classification error) comparisons. The different colors represent the different methods for analyses. The horizontal dash line is the lowest value corresponding to the best method.

### Independent Validation Results: Cross-dataset Performance

Although the six-fold cross-validation experiments above provide useful information, imputation will be performed in dataset(s) distinct from the one based on which training models are built. To provide more valid performance estimates and to assess the utility of models trained by CUE and by the individual methods, we examined their performances across the two datasets. With the presence of many systematic differences between the two datasets, we first attempted to correct for batch effects. Specifically, Combat ^36^ was used to generate a harmonized dataset after pooling together data from the two cohorts. Methylation prediction models were then trained with the harmonized ELGAN dataset and tested on the harmonized PTSD dataset. Among six imputation methods, CUE achieved the highest accuracy (95.48%) and the lowest predicted RMSE (0.0704). KNN was the fastest model with ∼1.35% loss in accuracy and 0.0189 loss in RMSE (Table 2). While this cross-dataset comparison is complicated by the different tissues in which methylation was assessed, these comparisons demonstrate the superior performance of the CUE method.

**Table 2.** Cross-dataset Imputation Performance.

| **Train: ELGAN**<br>**Test: PTSD** | **Test Performance** |  |  |
| --- | --- | --- | --- |
|  | **Accuracy** | **RMSE** | **Time** |
| **KNN** | 94.13% | 0.0893 | **1.5hrs** |
| **Logistic** | 92.30% | 0.2548 | 1.5hrs |
| **PFR** | 91.28% | 0.1341 | 5hrs |
| **RF** | 95.30% | 0.0706 | 1.5hrs |
| **XGBoost** | 94.90% | 0.0785 | 1.5hrs |
| **CUE** | **95.48%** | **0.0704** | 11hrs |

### DNA Methylation Varies across Different Tissue

DNA methylation data varies across different tissues inherently ^32, 37^. We classified DNA methylation probes into three categories: “unmethylated” if the *β* values across all samples in the cohort are less than 0.5; “methylated” if the *β* values across all samples in the cohort are greater than 0.5; “bilateral” otherwise. The proportions of three different categories of probes were reported for 339,033 HM850-targeted probes (**Figure 3**). For the PTSD cohort, almost 91% of probes were either methylated or unmethylated, while this is true of only 62% of probes in the ELGAN study. In general, the bilateral probes were more difficult to impute because of their inherent complexity, including true bilateral distribution patterns in some cases. One indicator of this complexity is that variance for these probes tended to be larger than the rest of the tested probes.

**Figure 3.**
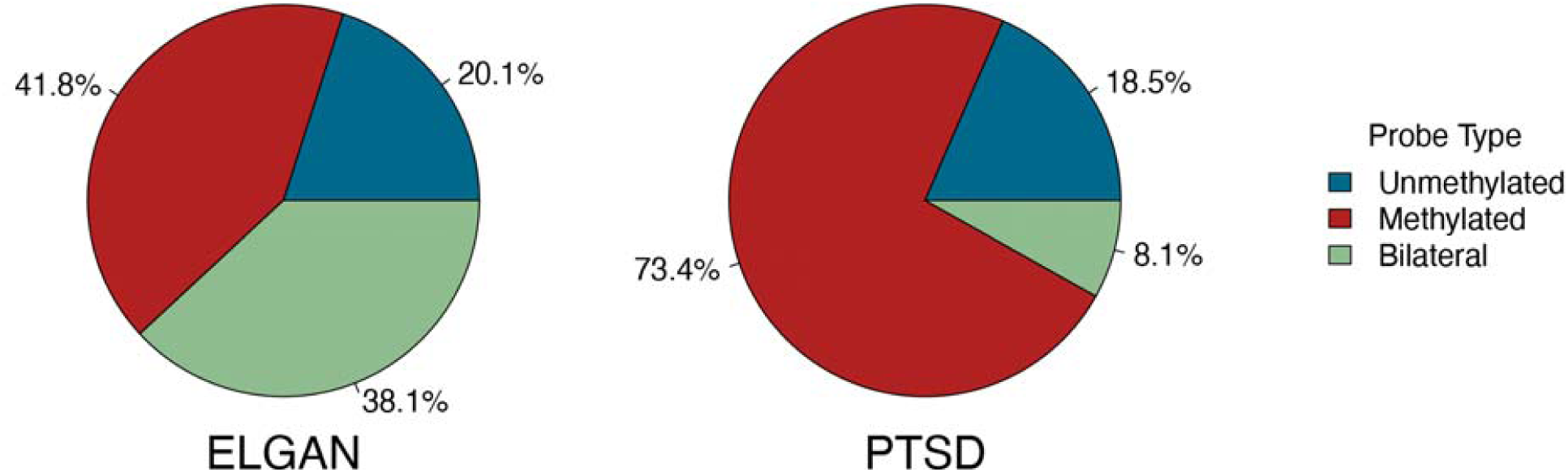
Proportions of three-category HM850-specific probes in PTSD and ELGAN.

## Discussion

In this study, we developed an ensemble method, CUE, to enhance prediction accuracy when imputing methylation between the HM450 and HM850 BeadChip platforms. Our initial goal was to extend our previously developed PFR framework ^29^ and systematically evaluate its performance with multiple alternative methods. However, our results suggested that there is no single DNA methylation imputation method available that performs best across different datasets or tissues, which motivated the development of the CUE method. Under three different scenarios (cross validation within PTSD dataset, cross validation within ELGAN dataset, and cross cohort imputation) with data from two different cohorts (one with samples from placenta and the other with samples from whole blood), CUE outperformed all five imputation methods in terms of both predicted RMSE (measuring methylation values with a continuous scale) and predicted accuracy (dichotomizing methylation values). CUE also led to a larger number of probes that passed post-imputation quality control than any other imputation method tested.

CUE produced accurate imputation results when the training and test data characterized the same tissue under similar conditions. With the CUE imputation framework, data can be combined from multiple methylation platforms, enabling higher resolution and more powerful downstream analysis, as long as an appropriate reference panel including samples assayed from both platforms is available. For example, the combined dataset can be used to boost the power of not only EWAS studies, but also DNA methylation quantitative-trait loci (mQTL) analyses, integrative multi-omics analyses, and other applications. Although in theory imputed and directly assessed methylation values could be combined directly, if possible, we recommend meta-analysis over mega-analysis to guard against potential batch effects from pooling together imputed and experimentally measured methylation levels. Regardless of the epigenetic architecture underlying phenotype(s) of interest, it is anticipated that this method will facilitate more efficient utilization of methylation data from multiple platforms and foster advances in understanding the role of DNA methylation on phenotype(s) of interest.

Despite the overall high imputation accuracy, quality control after imputation further improves imputation performance. As shown in Figure 3, the predicted RMSE of all the tools were reduced by 37.9%-50.0% after post-imputation QC in PTSD. For different datasets (tissues), we selected different quality control criteria (see more details in supplementary materials) according to cross-validation metrics within each dataset. We recommend similar basic QC filtering for all users of our CUE method, but users are free to adjust their QC criteria through our R package to find the best fit for their tissue and study.

DNA methylation data inherently vary across tissues, based on our results and those of others ^32, 37^, suggesting that it would be prudent to train separate imputation prediction models for different tissues. From this study, we provide two sets of imputation models: one for whole blood and the other for placenta. Investigators can therefore complete their own imputation of placental or whole blood HM850 CpG sites using their own HM450 data, without access to their own reference panel. Our method is also easily useable for imputation in other tissues, provided the user can supply a reference dataset assayed on both HM450 and HM850.

In summary, findings in this study suggest that the CUE ensemble methylation imputation method is valuable for imputing from HM450 to HM850. We hypothesize that CUE may also be helpful as new methylation arrays continue to be developed. This study is the first to impute from HM450 to HM850 using data from two different tissues. Using information at 248,421 HM450 CpG sites (sites overlapping between ELGAN and PTSD, and without missingness in our samples), CUE was able to accurately impute 289,604 HM850 sites in the PTSD whole blood samples and 238,090 sites in the ELGAN placenta samples. It is anticipated that the CUE method as well as the pre-trained imputation models across the two tissues will be of value to many investigators, facilitating more powerful epigenetics studies with either HM450 data or a mixture of HM450 and HM850 data. Ensemble methods like CUE may also benefit in more general settings for other tissues and with other future methylation assessment platforms.

## Methods

### Study Sample

Data were used from both the ELGAN study and the PTSD study. Samples from the PTSD genetics repository were from the Translational Research Center for TBI and Stress Disorders (TRACTS), a VA Rehabilitation Research and Development National Center for TBI Research at VA Boston Healthcare System. Informed consent was obtained from all PTSD subjects at the time of study inclusion. ELGAN study enrolled infants born < 28 weeks of gestion during 2002-2004, in five states and 14 hospitals in the United states ^35^. Detailed procedure regarding sample recruitment, main characteristics of study samples, and methylation measurements are presented in previous publications: Logue *et al*. for PTSD ^34^ and Santos *et al*. for ELGAN ^35^.

### Preprocessing of Methylation Data

The data used in this study have been pre-preprocessed previously. Logue *et al*. have previously published on the 145 samples from the PTSD cohort with methylation measured both by HM450 and HM850 ^34^. The cleaning and processing of this dataset according to a consortium-developed pipeline ^38^ has been described in detail elsewhere ^34^. Briefly, the PTSD dataset was first corrected for the individual-level background noise using GenomeStudio and then cleaned with the CpGassoc package and the ChAMP package in R ^39^. Detailed data cleaning and processing of the PTSD dataset was previously described ^34^. In this paper we further excluded 1 sample as it had a missing rate of >69% (568,833 missing probes among 820,611 probes) on the HM850 array, keeping 144 complete samples. Additionally, 127 subjects from ELGAN study ^35^ were selected based on the availability of placental samples with DNA methylation data assessed using both HM450 and HM850. DNA methylation data for the ELGAN dataset were first pre-processed by the minfi package ^40^. Then functional normalization was used for background subtraction and dye normalization. Santos *et al*. used the ComBat function from *sva* package to adjust for batch effects from two platforms, HM450 and HM850 ^36^. The detailed placenta tissue collection and other assessments of DNA methylation for the ELGAN dataset can be found in prior publications ^41-43^.

Since imputing sporadic missing data was not the focus of our work, probes with any missing values were removed. One could apply methods similar to those developed for gene expression data ^44-47^ to impute the sporadic missing values at directly assayed CpG sites. After removing sporadic missing values, probes were filtered to keep the common complete (no missing data) probes shared between two cohorts, which would make the assessments of different imputation models on two cohorts comparable. This left a total of 248,421 probes for HM450 and 587,454 probes for HM850 for the ELGAN and the PTSD cohorts respectively. A total of 248K HM450 probes were used as explanatory variables and the 339K HM850 specific probes were used as response variables.

Beta values (*β*s) were used to compare the imputation results. To ensure a Gaussian distribution ^48, 49^ for the PFR model, we employed M values, defined as *M* = logit(*β*) = log_2_ [*β*/(1 - *β*)], instead of the raw *β*s to impute and later transformed the imputed M values back (using the inverse of logit function) to *β* for comparisons. Again, *β* represents the intensity of the methylated probes over unmethylated probes.

### CUE: CpG impUtation Ensemble

The primary goal was to efficiently and accurately impute HM850-only probes (response variable) using probes from HM450 (explanatory variables). Specifically, for each target HM850 probe, we have *N* observations and for each observation or sample *i* = 1, 2, …, *N*, we have data [*Y*_*i*_, *Z*_*i*_], with *Y*_*i*_ as the transformed DNA methylation level at the target HM850 probe and Z_*i*_ is the *p*-dimensional vector of predictors (or covariates, explanatory variables; in this paper we use these three terms interchangeably).

In this work, we considered several imputation methods and benchmarked their performances on our three datasets across two cohorts. No single method was found to uniformly outperform the other methods for every CpG site tested and across different tissues or datasets. We therefore developed the following simple yet effective ensemble approach to boost prediction accuracy. Given *M* prediction estimators for the *j*-th CpG site, namely 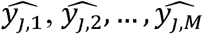, our ensemble prediction has the form 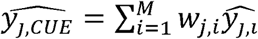 as an weighted estimation with weights w_*j, i*_ for i = {1,2, …, M}. Here we list three different approaches to select the weights. First, equal weights: 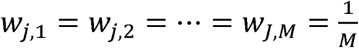; Second, best-single-method weights (0-1 weights): w_*j, best*_ = 1, the other weights =0; Third, theoretically-optimal weights: 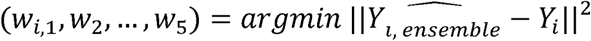. The equal-weights approach is simple and robust but could not guarantee the improvement of the performance. The best-single-method-weights approach and theoretically-optimal-weights approach would be guaranteed to be no worse than any single imputation tools by design. The theoretically-optimal-weights approach is actually the best linear fitting on the training data, which tends to overfit on the training data.

In this study, we adopted the best-single-method weights, seeking a balance between imputation performance and robustness. Based on the training results, we select the best method for imputation at each CpG site and employ that model for the final prediction. Here the model comparison criterion is out-of-bag predicted MSE. Suppose the *k*-th model outperforms other methods on a CpG site (i.e., with lowest out-of-bag prediction MSE), then *w*_*k*_ = 1 and *w*_*i*_ =0 for *i* ≠ *k*. Namely, 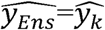 for this CpG site if the *k*-th model performs the best. Consequently, the performance of the ensemble method outperforms other single methods by design.

### Imputation Quality Assessment (Cross Validation)

Six-fold cross-validation was used to assess imputation quality. For each split, the full dataset was randomly divided into a training set, consisting of 5/6 of the total samples, and a testing set (1/6 of the total samples). For each testing set, only the data that were represented by probes common to HM450 and HM850 (shared probes, or predictor probes) were included, and masked methylation values of HM850-specific probes. For the training set, we employed methylation measurements on the shared probes as predictors to impute methylation values at HM850 specific probes. Since most HM450 probes are measured by both HM850 and HM450 platforms, the predictors used in our model can be methylation levels for these shared probes measured from either array. Note that our prediction model was built under the realistic and thus more challenging scenario where we used as predictors the measurements from HM450 array instead of those from HM850 array, which would require the training dataset had measurements from both arrays. Specifically, we first fitted our PFR model, learning the relationship between the methylation values of the shared and HM450-specific probes. Second, we used the fitted model to impute the masked values of HM850 probes from the HM450 data in the testing set. In the end, we evaluated the imputation performance by integrating imputation results from all 6 splits.

### Imputation Quality Measures and Quality Control Metrics

As measures of the imputation quality, we employ both the predicted RMSE and the accuracy with 0.5 as the threshold. Conventionally, if the raw methylation value was above the threshold, it was termed “methylated,” or “unmethylated” if below the threshold. Two quality control criteria were employed: for PTSD: the probe-level predicted RMSE < 0.05 and the probe-level predicted accuracy > 95% when dichotomizing DNA methylation level at a cutoff of 0.5; for ELGAN: the probe-level predicted RMSE < 0.1 and the probe-level predicted accuracy > 90%.

### Penalized Functional Regression (PFR) Model

Here we present a penalized functional regression model ^33^ with minor modifications. We also observe *X*_*i*_ (t), indexed by *T*_*i*_ =*t* representing sample-specific density function of the DNA methylation levels measured by HM450. Previous work has been shown that incorporating non-local density information improve the imputation accuracy when imputed form HM27 to HM450 ^323232^. We consider the following functional linear regression model: *Y* _*i*_ = *α* + ∫ X _*i*_ (t) *β* (t)dt + *Z*_*i*_*γ* + *ε*_*i*_, where *β* (t) ∈ ℒ^2^(*R*) characterizes the effect of density function *X*_*i*_ (t) when *T*_*i*_ = *t*. α is the grand mean and *γ* denotes the vector of regression coefficients corresponding to the vector of covariates *Z*_*i*_, 25 downstream probes and 25 upstream probes to each target probe as the local covariates.

Functional predictors *X* _*i*_ (t) were incorporated into the model to capture methylome-wide information, besides methylation levels from local probes encapsulated in *Z*_*i*_. According to the probe’s relative location to a CpG island, we first defined five groups: “CpG Island,” “North Shore,” “South Shore,” “North Shelf,” and “South Shelf” ^9^ (see **Figure 4**). The annotated group information can be found on Illumina’s website.

**Figure 4.**
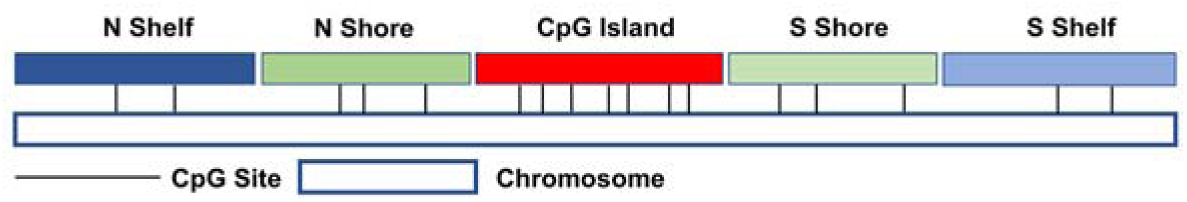
Five groups of CpG sites according to their relative location to CpG islands.

The DNA methylation function *X* _*i*_ (t) for a particular target probe was estimated via kernel density estimation using the DNA methylation data from all HM450 probes falling in the same group as the target probe. To perform model fitting, we projected the functional term *β* (*t*) onto a linear spline basis, then the model was reduced to a mixed effect model with *K*_*b*_ random effects, where *K* _*b*_ is the order of the linear basis. Previous studies have shown that the choice of the number of knots (the order for the linear basis and the knots for kernel density) is not important as long as it is large enough to capture the maximum complexity of the regression function ^32, 50, 51^. The advantage of the PFR approach is that it borrows information from the non-local probes. A limitation of PFR is that the run time of PFR is relatively long compared to other single imputation approach.

## Disclosure statement

No potential conflict of interest was reported by the authors.

## Funding

This work was supported by the National Institute of Health under grants including 5U01NS040069-05 (AL), 2R01NS040069-06A2 (KCK), UH3OD023348 (TMO and RCF), R01HD092374 (TMO and RCF), K23NR017898 (HS), T32HL129982 (LMR), 5R01HL129132-04 (YL) and from VA BLR&D Merit Award I01BX003477 (MWL).

## Notes

### Competing Interest Statement

The authors have declared no competing interest.

