## Supplemental File for "CUE: CpG impUtation Ensemble for DNA Methylation Levels Across the Human Methylation450 (HM450) and EPIC (HM850) BeadChip Platforms"

Gang Li
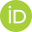
 ^a^, Laura Raffield
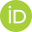
 ^b^, Mark Logue^c,d,l,m^, Mark W Miller
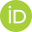
 ^c,d^, Hudson P. Santos Jr
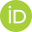
 ^g,h^, T.Michael O’Shea
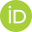
 ^i^, Rebecca C. Fry
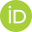
 ^h,j,k^, Yun Li
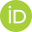
 ^b,e,f^

^a^Department of Statistics and Operations Research, University of North Carolina at Chapel Hill, Chapel Hill, NC, USA;

^b^Department of Genetics, University of North Carolina, Chapel Hill, NC, USA;

^c^National Center for PTSD at VA Boston Healthcare System, Boston, MA, USA;

^d^Department of Psychiatry, Boston University School of Medicine, Boston, MA, USA;

^e^Department of Biostatistics, Gillings School of Global Public Health, University of North Carolina, Chapel Hill, NC, USA;

^f^Department of Computer Science, University of North Carolina, Chapel Hill, NC, USA;

^g^School of Nursing, University of North Carolina, Chapel Hill, NC, USA;

^h^Institute for Environmental Health Solutions, Gillings School of Global Public Health, University of North Carolina, Chapel Hill, NC, USA;

^i^Department of Pediatrics, School of Medicine, University of North Carolina, Chapel Hill, NC, USA;

^j^Department of Environmental Sciences and Engineering, Gillings School of Global Public Health, University of North Carolina, Chapel Hill, NC, USA;

^k^Curriculum in Toxicology and Environmental Medicine, University of North Carolina, Chapel Hill, NC, USA;

^l^Biomedical Genetics, Boston University School of Medicine, Boston, MA, USA;

^m^Department of Biostatistics, Boston University School of Public Health, Boston, MA, USA.

**Table S1.** Numbers and Ratios of Well-imputed Sites. Quality control (QC) criterias for PTSD: RMSE<0.05 and Accruracy > 95%; QC for ELGAN: RMSE<0.1 and Accruracy > 90%. The ratios are out of 339,033 HM850-only CpG sites.

|  | PTSD (%) | ELGAN (%) |
| --- | --- | --- |
| KNN | 249,425 (73.6%) | 181,304 (53.5%) |
| Logistic | D.N.C. | 93,470 (27.6%) |
| PFR | 269,745 (79.6%) | 213,355 (62.9%) |
| RF | 249,425 (73.6%) | 236,645 (69.8%) |
| XGBoost | 285,330 (84.2%) | 224,317 (66.2%) |
| CUE | **289,604 (85.4%)** | **238,090 (70.2%)** |

**Fig S1.** Imputed performances for probes before (left, hatched) and after QC (right, no hatches) in ELGAN. (**a**) RMSE comparisons. (**b**) 1- Accuracy (classification error) comparisons. The different colors represent the different methods for analyses. The horizontal dash line is the lowest value corresponding to the best method (CUE) after QC.

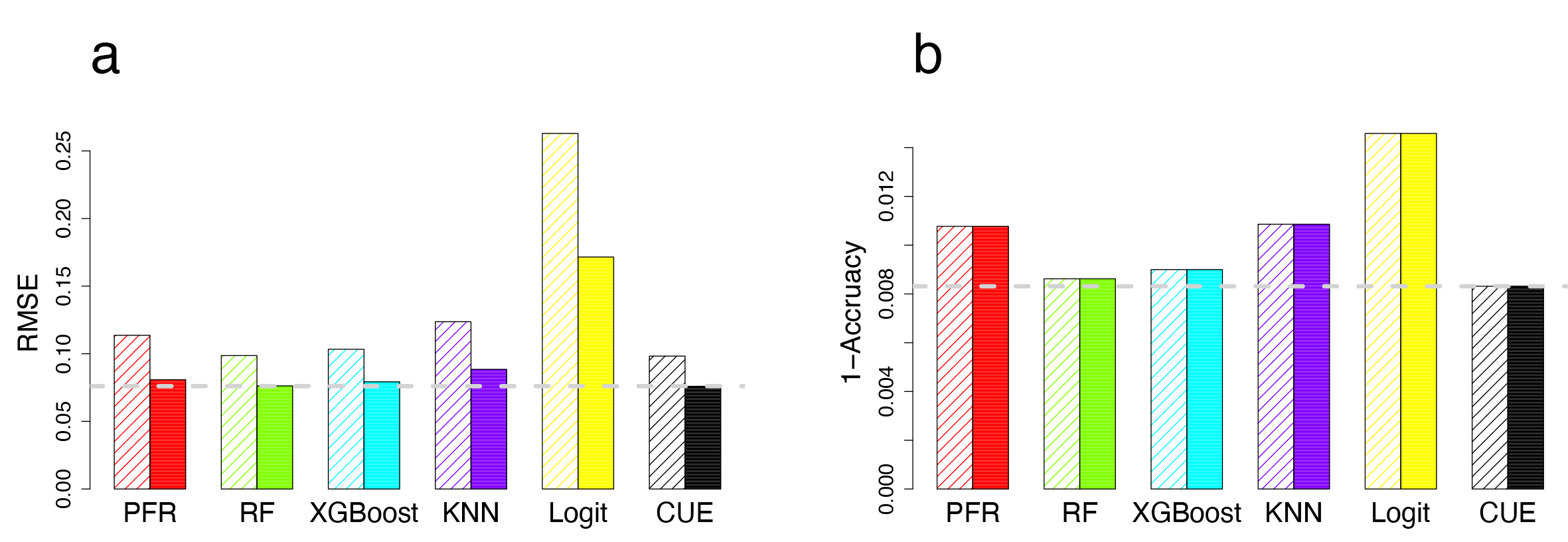
